# Coordinated Discordance: Strategic Nest Attendance for Chick Rearing in Monogamous Seabird

**DOI:** 10.1101/2025.04.26.650791

**Authors:** Daisuke Ochi, Nariko Oka, Yutaka Watanuki

**Affiliations:** Fisheries Resources Institute, Japan Fisheries Research and Education Agency; Yamashina Institute for Ornithology; Graduate School of Fisheries Science, Hokkaido University

**Keywords:** Bi-parental care, Chick provisioning, Nest attendance, Nest guard, Parental coordination, Streaked Shearwater

## Abstract

Behavioral coordination between mated pairs plays a critical role in the reproductive success of many species practicing bi-parental care. In pelagic seabirds such coordination may help cope with the challenges posed by spatially heterogeneous and limited food availability. This study investigated the relationship between parental coordination and reproductive performance by analyzing nest attendance patterns of breeding pairs of the Streaked Shearwater *Calonectris leucomelas* during the chick-rearing period on Mikura Island, Japan. Using an automated identification system, we recorded nest attendance and quantified coordination levels through the φ coefficient. Our analysis revealed that higher coordination levels were associated with reduced extended nest absence periods, potentially decreasing prolonged chick fasting. However, coordination levels showed no significant direct effect on chick growth rates, while regular provisioning frequency, which can be associated with the coordination, and meal size emerged as critical determinants of chick development. These findings indicate that, in Streaked Shearwaters, behavioral coordination during chick rearing contributes to indirect effect on chick development, and also, might reduce the duration of unguarded periods at the nest.

## INTRODUCTION

Behavioral coordination—where partners synchronize or alternate their actions—represents a sophisticated form of cooperation (Krause & Ruxton 2002). This coordination is particularly evident in parental care, where it can significantly impact offspring survival and parental fitness (Coulson 1966, Davis 1988). In long-lived species practicing biparental care, behavioral coordination often manifests as synchronized foraging and nest attendance (Coulson 1966, Niebuhr & McFarland 1983, Davis 1988, Booth et al. 2000). These species typically form long-term pair bonds, which may facilitate the development and refinement of coordinated behaviors over multiple breeding seasons.

In many species of offshore foraging seabirds, parents alternate long and short foraging trips (dual foraging strategy; Weimerskirch et al. 1994). This strategy serves distinct purposes: long foraging trips primarily allow parents to restore their own body condition by feeding in productive but distant foraging grounds, while short foraging trips maximize food delivery to chicks from nearby feeding areas, often at the expense of the parent’s condition (Weimerskirch 1998; Ochi et al. 2010). Recent studies on monogamous seabirds have reported coordinated discordance in nest attendance for chick provisioning, where one parent undertakes short foraging trips to compensate when the other parent is on a long foraging trip (Tyson et al. 2017). Similar coordinated provisioning patterns have subsequently been documented in various other seabird species (Kavelaars et al. 2021; McCully et al. 2022; Grissot et al. 2024).

Despite the growing interest in the coordinated provisioning behavior, there are significant challenges in quantifying the level of coordination. Many researchers have relied on subjective measures to indicate the level. For instance, foraging trip durations are often arbitrarily categorized into short and long trips using a fixed threshold, despite the fact that trip durations typically follow a continuous distribution (Ochi et al. 2016; Tyson et al. 2017). This makes it difficult to conduct comparative analyses. A more robust approach would involve using indicators that quantify the level of coordination as continuous variables rather than relying on arbitrary cut-offs. The development of such objective indicators would greatly contribute to our understanding of the significance of the coordinated behavior.

Previous studies have predominantly focused on the role of coordinated provisioning in maintaining consistent feeding rates and extended fasting period (Tyson et al. 2017; Ihle et al. 2019a). However, empirical support for this hypothesis has been inconsistent across studies. A more comprehensive evaluation of coordinated behavior should consider these multiple potential functions rather than focusing exclusively on feeding rates. By quantifying coordination using continuous indicators, we may develop a more detailed understanding of parental care strategies across diverse ecological contexts. For example, coordinated nest attendance patterns may make additional contributions to chick rearing by allowing parents to spend more time at the nest and thereby potentially deterring predators or conspecific intruders (Leniowski and Wegrzyn 2018; Ihle et al. 2019a).

Through this study, we examined a monogamous seabird species of shearwater to investigate differences in coordination behavior between pairs and their reproductive output. Our research consisted of three main components: (1) measurement of coordination levels using objective indicators applicable even when modes of foraging trip are not clearly distinguishable, (2) analyzing the relationship between coordination levels and both feeding and chick growth, and (3) examining the relationship between coordinated pairs and nest guarding behaviors.

## METHODS

Streaked Shearwaters *Calonectris leucomelas* represent an appropriate species for investigating coordinated nest attendance between mated pairs due to their distinctive life history traits and ecological constraints. Their strong reliance on biparental care, clear nest attendance patterns, and the necessity to conduct foraging trips of varying durations provide an excellent framework for examining coordination strategies. These shearwaters breed on Mikura Island (33°52’N, 139°14’E), Japan, which is the largest breeding colony of this species (Oka 2004). This species is monogamous, with both males and females sharing incubation and provisioning duties. They are burrow-nesting seabirds that lay a single egg per breeding attempt. Adults visit the colony exclusively at night to feed their chicks, and by morning, all parents had departed the island for foraging trips of varying durations. Although this breeding colony is situated at a considerable distance from highly productive foraging areas, suggesting a dual foraging strategy with long and short trips, the boundary between these trip types remains notably indistinct in this population (Ochi et al. 2016).

The field study was carried out from August to November 2005 on Mikura Island after receiving permission from the Tokyo Metropolitan Government, Japan (license no. 27). We captured both male and female parents of Streaked Shearwaters nesting in artificial nest boxes (62 × 48 cm; 35 cm high, with a plastic entrance tunnel 25 cm in diameter and 1–2 m long). We weighed each parent and placed a plastic band with an attached cylindrical magnet around its tarsus (described below). Handling of each bird was completed within 10 minutes. The sex of all birds was determined by their vocalizations during handling: calls of males are high-pitched, whereas those of females are low-pitched (Arima & Sugawa 2004).

Nest visits by the parents were recorded by an automatic attendance recording system (Ochi et al. 2006; 2010). Plastic bands (3 cm long; mass, <3 g; <1% of the adult body mass), to which a magnet (1 mm in diameter; 2.4 cm long, <50% of the length of an adult tarsus) was affixed by epoxy glue, were attached to the tarsi of all adults so that the magnetic polarity was oriented differently for male and female parents. Two Hall sensors (HW-300A. Asahi-Kasei Electronics Corp., Tokyo, Japan) were set under the entrance of each nest burrow. These sensors detected changes in the local magnetic fields induced by parent movements, recording precise entry and exit times when magnets passed through the burrow entrance. This system provided three key temporal metrics: arrival time, departure time, and nest stay duration. Signal changes were recorded at 1.5-s intervals with data loggers (AKI-Logger; Akizuki Denshi Tsusho, Saitama, Japan or HOBO H8; Onset Computer Corp., Bourne, MA, USA). During heavy rainfall, some Hall sensors accidentally shorted out due to water infiltration and stopped functioning; however, this did not affect the identification of returning individuals, except that they were no longer able to distinguish between arrivals and departures.

Chicks were weighed with a Pesola spring balance (Pesola AG, Baar, Switzerland) to the nearest 1 g in the morning (0600 hours) and evening (1800 hours) every day. The handling time of the chicks was minimized to no more than two minutes. We performed a linear regression using daily chick mass measurements and treated the regression coefficient as the chick growth rate (g/day). The provisioning mass was estimated based on the difference in chick weight (*DEF*) between one evening (*W*_18_) and the following evening (Provisioning mass = 1.102*DEF* + 0.113*W*_18_ – 0.498, Inoue et al. 2009). We also treated mean daily provisioning mass as provisioning rate.

The φ coefficient, a measure of association between two binary variables (i.e. present/absence), was calculated to quantify the strength of coordination in attendance patterns. This statistic, equivalent to Pearson’s correlation coefficient for 2×2 contingency tables (Cohen, 1988), is derived using the formula:

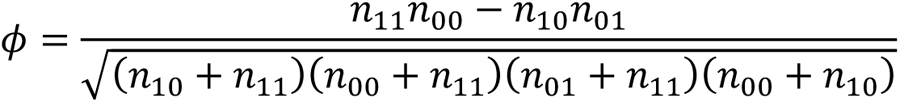

where n_11_ and n_00_ represent concordant attendance by pairs (both variables present/absent), while n_10_ and n_01_ denote discordant attendance. The denominator adjusts for marginal distributions, producing values between -1 (perfect discordant association) and 1 (perfect concordant association), with 0 indicating statistical independence.

To examine relationships between level of coordination (φ coefficient) and chick development, we calculated Pearson’s correlation coefficients to assess the strength and direction of associations between the following pairs of variables: (1) φ coefficient and proportion of extended nest absence, (2) proportion of extended nest absence and chick growth rate, (3) φ coefficient and chick growth rate, (4) φ coefficient and provisioning rate (g/day), and (5) provisioning rate and chick growth rate. We defined extended nest absence (= extended fasting) as periods exceeding three consecutive days without either parent present, based on the observation that long foraging trips to productive feeding grounds from this colony typically require an average of three days (Ochi et al. 2016). These extended absences likely represent situations where both parents simultaneously engaged in long trips.

We assumed that pairs might be coordinating their discordant attendance to improve nest guarding effort. To distinguish different coordination strategies, we attempted to categorize pairs into two groups—those exhibiting coordinated attendance and those that did not—using the φ coefficient as our classification metric. However, given the non-random nature of the attendance patterns, we needed a more practical approach than the standard χ^2^ test to determine significance. For instance, extremes such as daily returns to the nest or never returning at all are highly unlikely in nature.

To address this, we employed simulation-based validation. By generating random attendance patterns based on the observed foraging trip duration data and comparing the simulated φ coefficients with the actual φ coefficients, we could assess whether the observed pattern represented intentional behavioral coordination rather than chance alignment of independent foraging schedules. This method allowed us to account for the inherent biases in the attendance patterns and provide a more accurate assessment of coordinated behavior among pairs.

To simulate attendance patterns, we used the observed trip duration patterns and observation periods for each individual in a pair as input data. The simulation process involved randomly sampling trip durations with replacement from the observed patterns to generate new trip sequences. These sequences were extended until their total duration exceeded the observation period plus a 20-day warm-up period. From these simulated trip sequences, we created attendance patterns, discarding the first 20 days as a warm-up and retaining the subsequent days corresponding to the actual observation period. This warm-up period was introduced to avoid the issue where the first day of all simulations is inevitably set as an “attend” by default. We then combined the simulated attendance patterns for each pair to create nest attendance patterns. Finally, we computed the φ coefficient for each simulated pattern. This entire process was repeated 10,000 times to generate a distribution of simulated φ coefficients for comparison with the observed values. Statistical significance was assessed by comparing the actual φ coefficient with the simulated distribution—observed values falling below the 5th percentile of the simulated distribution (i.e., lower than 95% of simulated values) were considered statistically significant under a one-tailed assessment. The simulation was implemented by R 4.4.2 (The R development core team) and the simulation code is included in Supplemental material S1.

We analyzed differences in arrival times, departure times and nest stay periods between coordinated and non-coordinated pairs. For all variables, we ran linear models, with pair coordination type (“coordinated” vs. “non-coordinated”) and individual sex as independent variables.

We set the level of statistical significance at *P* < 0.05 and represented measured and calculated data as means ± SD. All the statistical analyses were conducted using R 4.4.2 (The R development core team).

## RESULTS

The recording period at each nest ranged from 22 to 70 days. We recorded nest attendance when the chicks were between 5 and 82 days old. We used data collected before the chicks attained their maximum mass (62–82 days) because parents commonly cease provisioning to chick after that time (Oka et al. 2002). Trip duration ranged from 1 to 13 days (mean = 2.7± 2.6 days, N = 618; Fig. 1a). The nest absence period ranged from 1 to 9 days (1.52 ± 1.13 days; N = 504, Fig. 1b) among the observed pairs.

**Figure 1.**
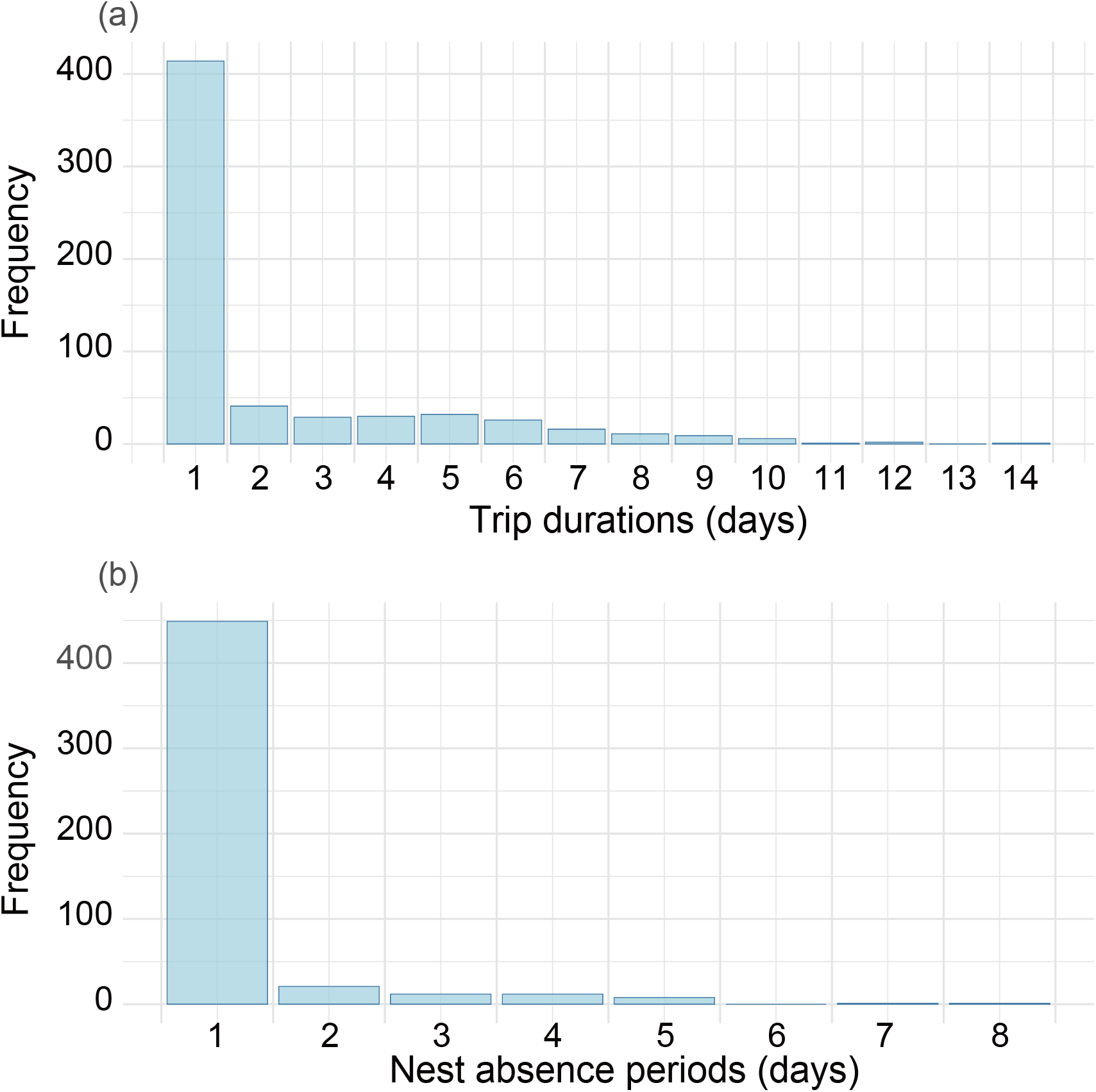
Frequency distributions of (a) foraging trip durations and (b) consecutive nest absence periods by pairs.

The chick at one nest showed notably delayed development, reaching a peak mass at 80 days of age without reaching any maximum equilibrium between the middle and the late nestling periods reported in previous studies (Oka et al. 2002). This was due to a male that nearly neglected chick rearing, very few nest visits during the second half of the chick-rearing period, rather than simply being uncoordinated with its mate. Another nest where a chick was predated by a feral cat before reaching maximum mass was excluded from growth rate analyses (but retained for φ calculation and analysis for nest guarding), as growth rates naturally decelerate approaching peak mass.

Based on the nest attendance patterns of 17 recorded pairs, we calculated φ coefficients and found that 16 out of 17 pairs exhibited negative values (discordant association; Fig. 2). For clarity in subsequent analyses, we reassigned pair IDs in ascending order of φ coefficients. The attendance patterns of the two pairs with the highest and the two pairs with the lowest φ coefficients are shown in Fig. 3.

**Figure 2.**
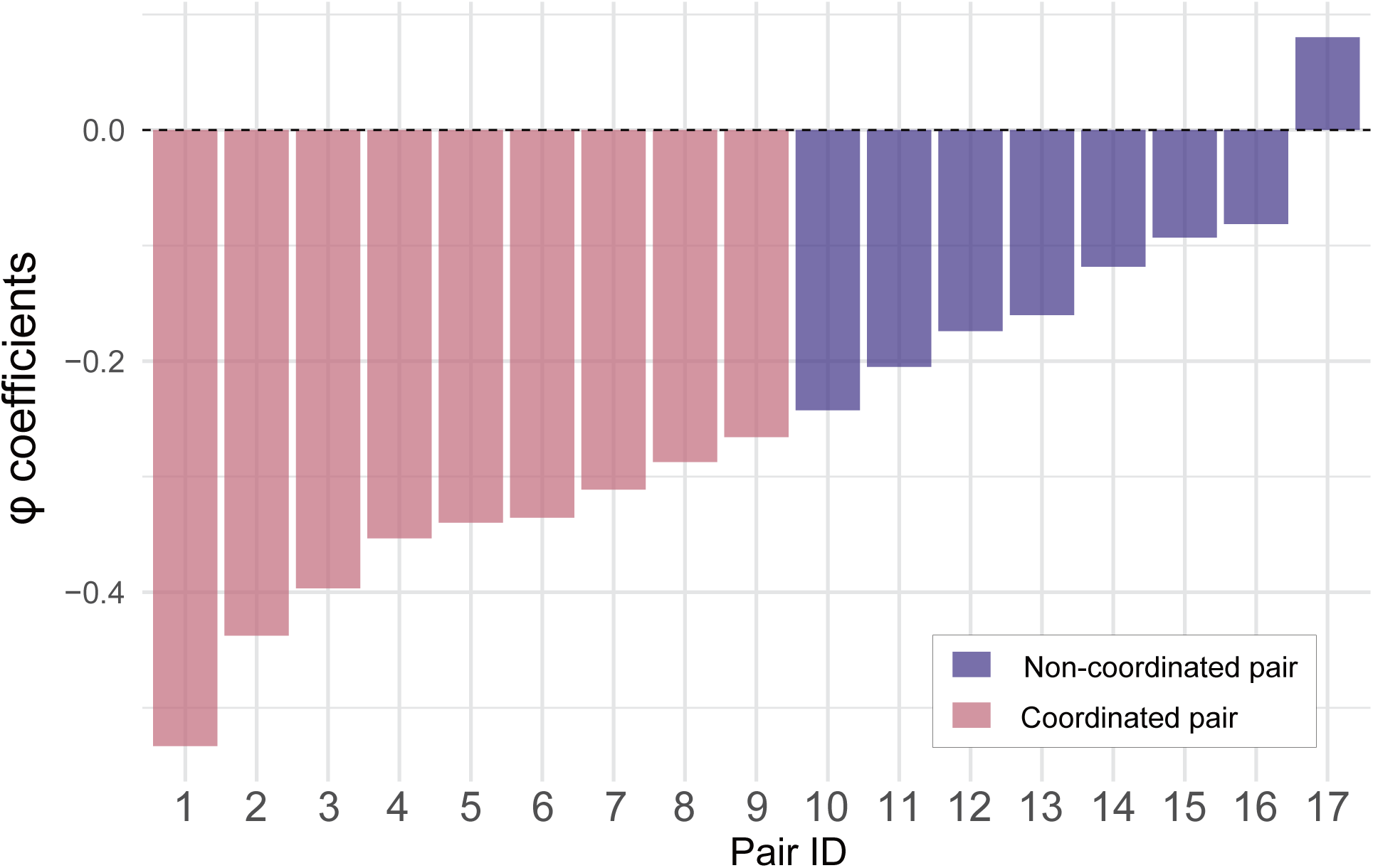
Coordination levels (φ coefficients) calculated from observed pair attendance patterns. Pink-filled bars represent pairs categorized as “coordinated” based on resampling simulations, while purple-filled bars indicate pairs categorized as “non-coordinated”.

**Figure 3.**
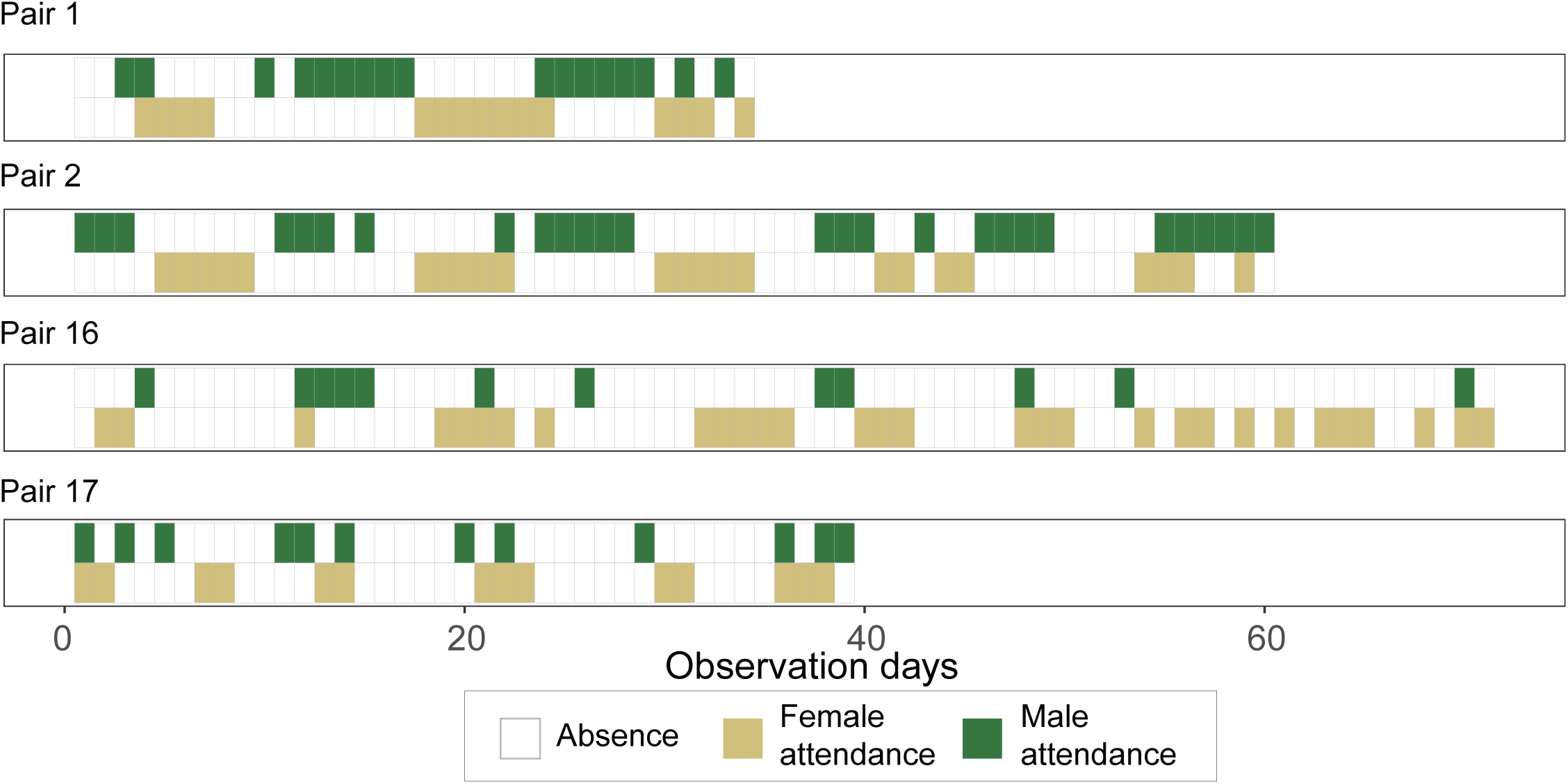
Nest attendance patterns of two pairs with the highest φ coefficients and two pairs with the lowest φ coefficients, as recorded by the automated detection system.

All survived chicks grew normally (mean growth rate: 6.7 ± 2.2 g/day, *N* = 16) and estimated provisioning was 44.6 ± 7.4 g/day on average (N = 16). The φ coefficient showed a positive correlation with the proportion of extended nest absence (R = 0.55, p = 0.026; Fig. 4a), and there was a negative correlation between the proportion of extended nest absence and chick growth rate (R = − 0.54, p = 0.029; Fig. 4b). However, no significant correlation was observed between the φ coefficient and chick growth rate (R = − 0.16, p = 0.56). Meanwhile, provisioning rate demonstrated a strong positive correlation with chick growth (R = 0.73, p = 0.001; Fig. 4c) and no significant correlation between φ coefficient and provisioning rate (R = − 0.46, p = 0.08).

**Figure 4.**
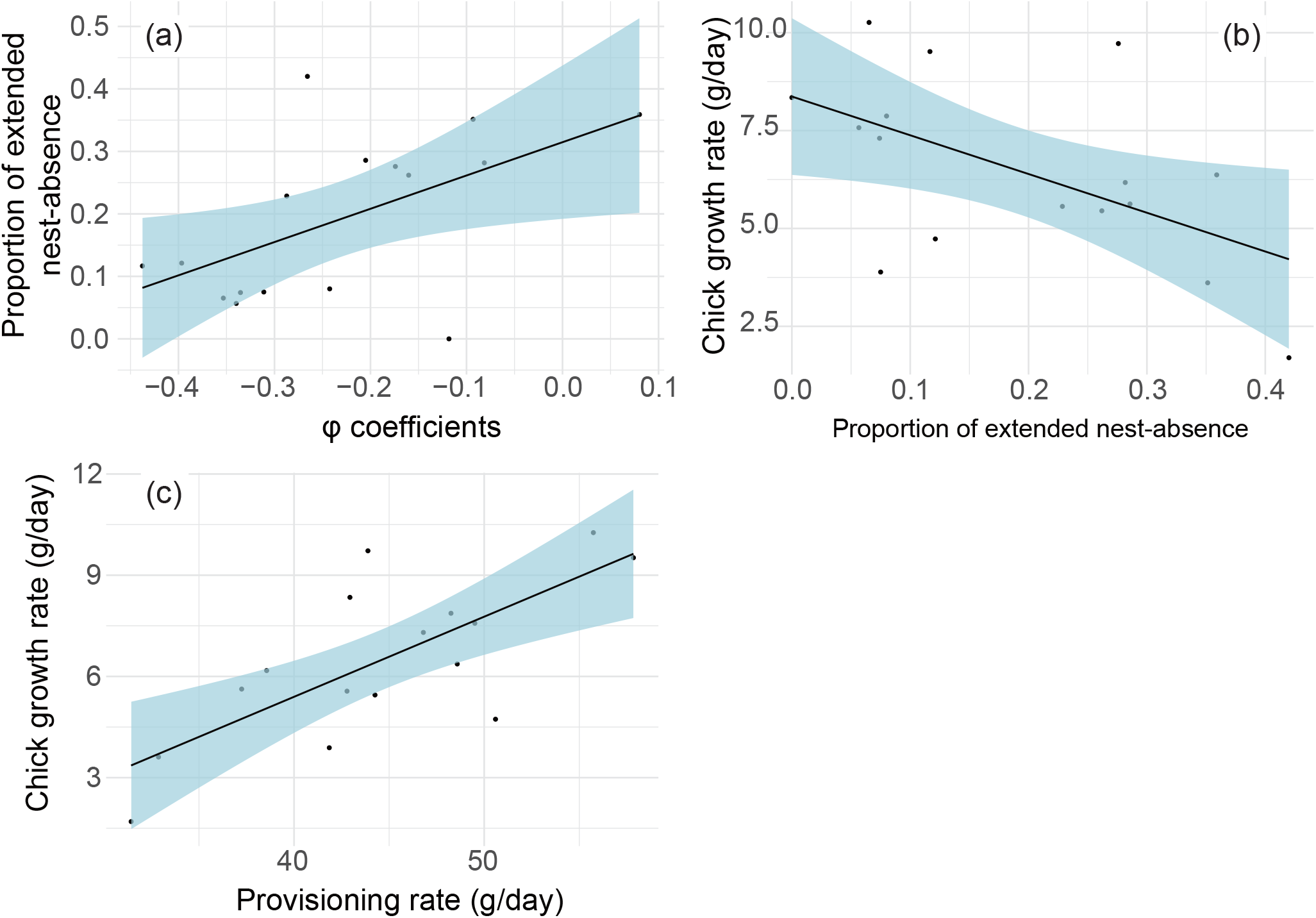
Correlations between φ coefficient, proportion of extended nest absences (>3 days), provisioning rate, and chick growth rate. (a) φ coefficient vs. proportion of extended nest absences, (b) proportion of extended nest absences vs. chick growth rate, and (c) provisioning rate vs. chick growth rate. Solid lines represent linear regressions, and shaded areas indicate 95% confidence intervals.

By comparing the distribution of φ coefficients obtained from attendance pattern simulations with the actual observed φ coefficients, we found that pairs with IDs 1–9 had observed values falling below the 5th percentile of the simulated distribution (Supplemental material S2). Based on this criterion, we categorized these pairs as “coordinated” pairs, while designating the remaining nine pairs as “non-coordinated” pairs.

The mean arrival time at the nests was 21:27 ± 2:47 (N = 557), with a mean departure time of 2:12 ± 2:25 (N = 556), and an average nest stay duration of 4hr45min ± 2hr58min (N = 556). Males arrived significantly later than females, while there was no significant difference between coordinated and non-coordinated pairs in arrival timing (Table 1, Fig. 5a). In contrast, departure times showed no sex difference, but individuals from coordinated pairs remained at the nest significantly later (Table 1, Fig. 5b). Consequently, individuals from coordinated pairs had significantly longer nest stay durations compared to those from non-coordinated pairs (Table 1, Fig. 5c).

**Table 1.** Summary of linear models testing the effects of pair coordination status and sex on nest arrival time, departure time, and nest resident duration.

| Variables | Coefficients | Std.<br>error | t | p |
| --- | --- | --- | --- | --- |
| **Arrival time** |  |  |  |  |
| Intercept | 21.10 | 0.21 | 100.42 | <0.001 |
| Sex<br>(Female as reference) | 0.57 | 0.24 | 2.45 | 0.02 |
| Pair type<br>(Non-coordinated pair as reference) | 0.13 | 0.24 | 0.54 | 0.59 |
| **Departure time** |  |  |  |  |
| Intercept | 25.49 | 0.18 | 142.46 | <0.001 |
| Sex<br>(Female as reference) | 0.24 | 0.20 | 1.19 | 0.24 |
| Pair type<br>(Non-coordinated pair as reference) | 1.06 | 0.20 | 5.27 | <0.001 |
| **Nest stay period** |  |  |  |  |
| Intercept | 4.39 | 0.22 | 19.74 | <0.001 |
| Sex<br>(Female as reference) | -0.33 | 0.25 | -1.34 | 0.18 |
| Pair type<br>(Non-coordinated pair as reference) | 0.93 | 0.25 | 3.72 | <0.001 |

**Figure 5.**
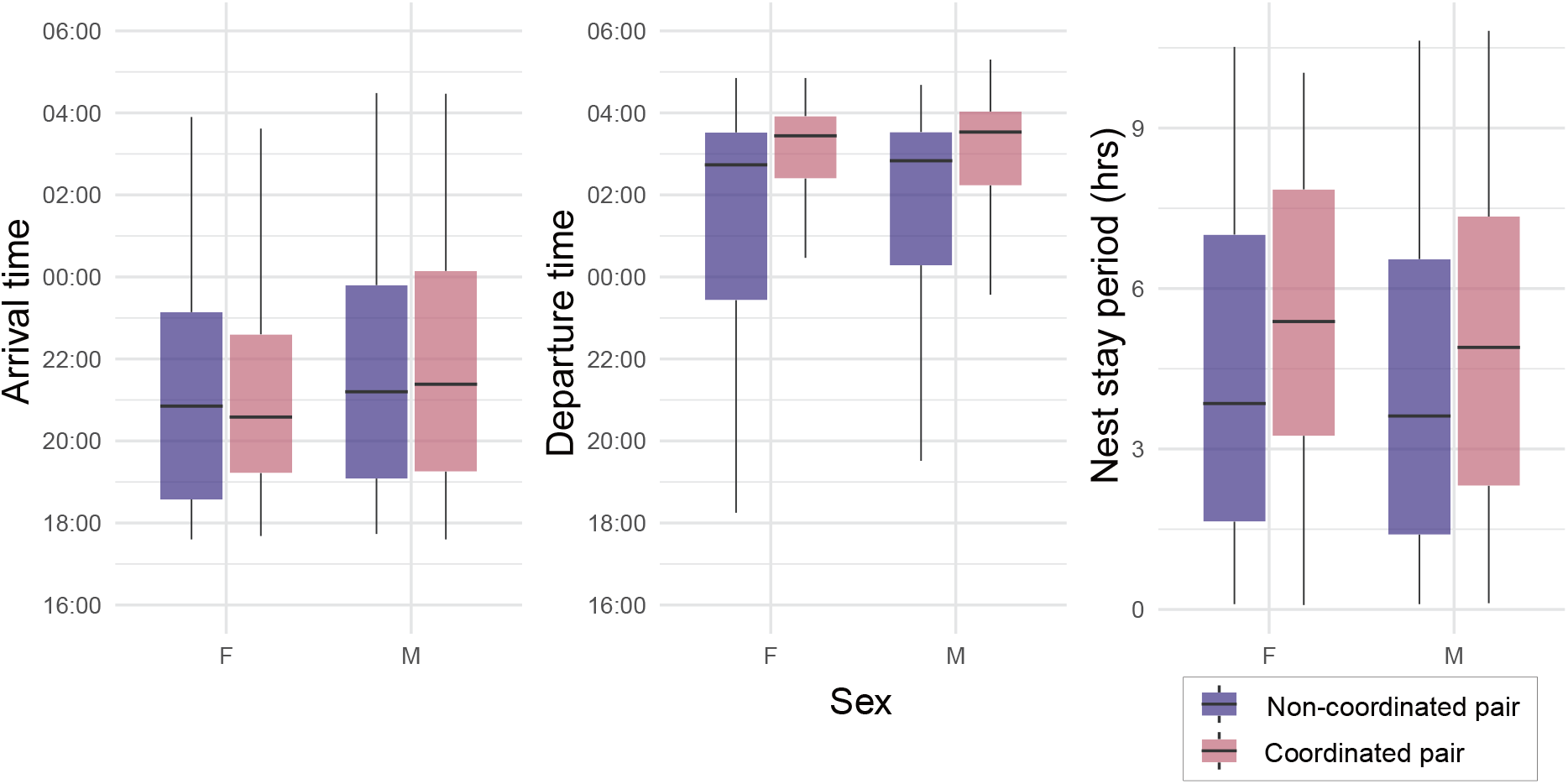
Box plots comparing (a) arrival times, (b) departure times, and (c) nest stay periods between coordinated and non-coordinated pairs, separated by sex.

## DISCUSSION

We employed the φ coefficient as an objective indicator to demonstrate differences in the level of coordination behavior between pairs. Our findings on pairs of Streaked Shearwater showed that the level of coordination itself did not show a significant relationship with chick growth. Interestingly, coordinated pairs remained at the nest until just before dawn and stayed in the nest for longer durations compared to non-coordinated pairs. Taken together, these findings suggest that their coordinated attendance behavior likely showed multiple aspects.

Level of coordination did not directly influence chick mass growth performance, supporting the findings of some previous studies (Wojczulanis-Jakubas et al. 2018; Ihle et al. 2019a; Grissot et al. 2019). Other research has suggested that coordination might only affect early chick growth (Tyson et al. 2017). As our study focused on middle to late-stage chicks, we were unable to examine this possibility. Providing larger amounts of food at higher frequencies (i.e., higher provisioning rate) on average appears to be effective for chick growth, rather than consistent provisioning at shorter intervals through coordinated attendance as we had hypothesized. If maximizing average provisioning rate were the primary objective, parents could simply maximize average provisioning rate by increasing feeding frequency or meal size on average regardless of the variances based on individual capacity (Ricklefs 1968) without needing to coordinate return timing.

Nevertheless, since extended fasting periods showed a negative correlation with growth rate, coordinated attendance may still have an indirect influence on chick development by reducing the frequency of prolonged fasting episodes. Extended fasting can negatively impact aspects beyond growth rate, including immune function, visual capabilities, and final body size, as indicated in previous studies (Ohlsson & Smith 2001; Fargallo et al. 2002; Kitaysky et al. 2005). Parents may coordinate their attendance to prevent these adverse effects, even if immediate growth benefits are not apparent. The observational approach of our research makes it difficult to completely separate these potential effects of feeding frequency from coordination levels; experimental manipulations, such as adjusting provisioning amounts, would be necessary for more definitive conclusions. A delayed chick growth was recorded only in the case that one of the parents nearly ceased provisioning to the chick in the later chick-rearing season (Pair 16 in Fig. 3). This case underscores the importance of consistent bi-parental care and suggests that a lack of parental cooperation can have measurable negative impacts on chick development.

The coordinated nest attendance may increase the time at nest hence may benefit the defense from conspecific interference, and maintenance of the nest burrow. Mikura Island, the breeding site, hosts both native predators such as Japanese Striped Snakes *Elaphe quadrivirgata* and Large-billed Crow *Corvus macrorhynchos*, as well as introduced predators including feral cats, Black Rats *Rattus rattus*, and Brown Rats *Rattus norvegicus* (Azumi et al. 2019, 2021). All of these except crows can approach the burrows during the night, making parental attendance at night potentially highly beneficial. However, the presence of invasive predators, feral cats, may create an evolutionary mismatch-while extended nest stays could enhance chick protection, they may simultaneously increase predation risk for attending adults, as these novel predators are not part of the species’ evolutionary history. Indeed, during our observations, we found one chick predation by a feral cat (evidenced by a decapitated carcass remaining outside the burrow; Nagata et al. 2022). This supports the considerable benefits of coordinated nest defense. Hostile intra-species interference may be another factor as suggested in Pelicans (Daigre et al. 2012) and in petrels (Dilley et al. 2019). Indeed, in Streaked Shearwaters, video observations have documented non-paired adults frequently entering nests and interfering with chicks (Inoue, Y. personal communication). While our study provides indirect evidence for nest guarding benefits through increased nest attendance duration, future investigations should incorporate direct metrics of nest guarding efficacy. It should be noted that invasive predators such as feral cats may introduce a novel risk, as their presence could negate some benefits of coordinated attendance by threatening both chicks and attending adults.

The φ coefficient utilized in this study can determine correlations between two time-series binary datasets (such as nest presence/absence), making it a highly versatile and simple indicator that could be readily applied to pairs of other animal species. For example, in species that return to the nest multiple times daily for feeding, coordination levels could be analyzed using the φ coefficient by setting time-block units of appropriate length for that species. The φ coefficient framework developed here enables cross-species or cross-population comparisons of coordination strategies as this approach can be applicable to the data with various biases.

While we have demonstrated coordination among pairs, the question remains: how do they adjust their timing? When exhibiting coordinated but discordant attendance patterns, pairs have few opportunities to meet directly, seemingly leaving no method for timing adjustment. Studies of Manx Shearwaters *Puffinus puffinus* suggest that communication may occur during moments when both parents simultaneously return to the nest (Tyson et al. 2017). In our study, even the highly coordinated pairs had nights when both partners were present simultaneously, providing potential opportunities for communication (Pair 1 & 2; Fig. 3). Additionally, coordinated pairs spent longer periods in the nest together, suggesting they may engage in extended communication when their visits overlap. Indirect communication through the response of chick is another possible mechanism. Reports on Streaked Shearwater indicate that chick condition and begging intensity can influence the timing of the return of mates (Ochi et al. 2010; Ogawa et al. 2015), though other theoretical research has proposed the environmental factors likely influence timing of arrival (Ihle et al. 2019b). In Streaked Shearwaters, adjustments in return timing based on weather conditions such as wind speed, have been also suggested (Van Tatenhove et al. 2018). A comprehensive investigation considering both individual circumstances and environmental factors would be necessary to fully understand this coordination mechanism.

## ACKNOWLEDGMENTS

We express our gratitude to Y. Inoue, K. Matsumoto, T. Deguchi, R. Yamashita, T. Tokunaga, H. Fujii, K. Hirata, Y. Watanabe and T. Yamamoto for their assistance during fieldwork and logistical support. We also thank K. Kazama, M. Ito, A. Takahashi, K. Matsumoto and Y. Iwata for their valuable comments during manuscript preparation. We appreciate the various suggestions provided by members of the Animal Ecology Laboratory, Faculty of Agriculture, and the Marine Ecology Laboratory, Faculty of Fisheries, Hokkaido University, which greatly contributed to the completion of this research. We also appreciate the support for providing the Hall sensors for the detection system by the Asahi-Kasei Electronics Corp. Finally, we are deeply indebted to the late S. Kurimoto for securing the research field, providing material support, and offering companionship during solitary fieldwork, among numerous other contributions. We extend our sincere appreciation to all those mentioned.

## AUTHORSHIP CONTRIBUTION STATEMENT

Daisuke Ochi: Conceptualization, Investigation, Methodology, Writing – original draft, Writing – review & editing. Nariko Oka: Funding acquisition, Writing – review & editing. Yutaka Watanuki: Funding acquisition, Supervision, Writing – review & editing.

## SUPPLEMENTAL MATERIALS

This study includes two supplemental materials:

Supplemental Material 1 (S1): R code for performing the φ coefficient resampling simulation based on observed trip duration patterns of breeding pairs.

Supplemental Material 2 (S2): Results of the resampling simulation, presenting the bootstrapped distribution of φ coefficients and their comparison to observed values.

