## Supplemental material S1 for "Coordinated Discordance: Strategic Nest Attendance for Chick Rearing in Monogamous Seabird"

```
# Supplemental material S1 Resampling Simulation Code ####
#
# Program to generate new trip patterns from trip length distributions obtained from observations
for each pair,?A
# create nest attendance patterns, combine them, and calculate the phi coefficient

library(tidyverse)
library(furrr)
library(psych)

# Setting for parallel processing
plan(multisession)

# Sample data ####
nests_df <- tibble(
  pair_id = as.character(1:5),
  observation_period = c(30, 28, 31, 30, 29),
  male_trip_lengths = list(
    c(2,1,3,2,1),
    c(3,2,1,4),
    c(2,2,3,1,2),
    c(4,1,2,3),
    c(2,3,1,2,1)
  ),
  femaletrip_lengths = list(
    c(2,3,1,2),
    c(1,3,2,2),
    c(3,1,2,3,1),
    c(2,2,1,3),
    c(1,2,3,1)
  )
)

# Define simulation functions ####
# 1. Resample from observed trip lengths to generate new trip sequences
# 2. Convert generated trip patterns to nest attendance patterns (binary data, daily)
simulate_patterns <- function(trip_lengths, observation_period) {
  # Function to generate trip pattern
  # The first day will always have nest attendance = 1, so we add 20 extra days as a warm-up period
  generate_trip_pattern <- function(trip_lengths, observation_period) {
    total_days <- 0
    generated_pattern <- c()
```

```
    while (total_days <= observation_period + 20) {
      trip <- sample(trip_lengths, 1, replace = TRUE)
      generated_pattern <- c(generated_pattern, trip)
      total_days <- total_days + trip
    }
    return(generated_pattern)
  }

  # Function to generate nest attendance pattern
  create_nesting_pattern <- function(trip_pattern, observation_period) {
    nesting_pattern <- unlist(map(trip_pattern, ~ c(rep(0, .x - 1), 1)))
    nesting_pattern <- nesting_pattern[1:(observation_period + 20)]
    return(nesting_pattern[-(1:20)])
  }

  nesting_counts <- length(trip_lengths)

  # Generate trip pattern and create nest attendance patter
  trip_pattern <- generate_trip_pattern(trip_lengths, observation_period)
  nesting_pattern <- create_nesting_pattern(trip_pattern, observation_period)

  return(list(trip_pattern = trip_pattern, nesting_pattern = nesting_pattern))
}

# Function to calculate phi coefficient based on contingency table
calculate_phi_coefficient <- function(pattern1, pattern2) {
  # Create contingency table
  tbl <- table(pattern1, pattern2)

  # Check if it's a 2x2 table
  if (all(dim(tbl) == c(2, 2))) {
    # Calculate phi coefficient
    return(phi(tbl))
  } else {
    # Return NA if not a 2x2 table
    return(NA)
  }
}

# Function to calculate number of nest attendances
calculate_nesting_counts <- function(nesting_pattern) {
  return(sum(nesting_pattern))
```

```
}

# Function to repeat simulation
run_simulation <- function(pair_id, male_trip_lengths, female_trip_lengths, observation_period,
iteration) {
  # Prepare male and female data
  male_data <- tibble(
    individual = "male",
    trip_lengths = list(male_trip_lengths),
    observation_period = observation_period
  )

  female_data <- tibble(
    individual = "female",
    trip_lengths = list(female_trip_lengths),
    observation_period = observation_period
  )

  # Run simulation
  male_results <- male_data %>%
    mutate(simulation = future_map2(
      trip_lengths,
      observation_period,
      ~ simulate_patterns(.x, .y),
      .options = furrr_options(seed = TRUE)
    )) %>%
    mutate(nesting_patterns = map(simulation, "nesting_pattern"))

  female_results <- female_data %>%
    mutate(simulation = future_map2(
      trip_lengths,
      observation_period,
      ~ simulate_patterns(.x, .y),
      .options = furrr_options(seed = TRUE)
    )) %>%
    mutate(nesting_patterns = map(simulation, "nesting_pattern"))

  # Generate pair nest attendance pattern
  pair_nesting_pattern <- map2(
    male_results$nesting_patterns[[1]],
    female_results$nesting_patterns[[1]],
    ~ as.numeric(.x | .y)
```

```
  ) %>% unlist()

  # Calculate phi coefficient
  phi_coefficient <- calculate_phi_coefficient(male_results$nesting_patterns[[1]],
female_results$nesting_patterns[[1]])

  # Calculate nest attendance counts from individual attendance patterns
  male_nesting_count <- calculate_nesting_counts(male_results$nesting_patterns[[1]])
  female_nesting_count <- calculate_nesting_counts(female_results$nesting_patterns[[1]])

  return(list(
    pair_id = pair_id,  # Return pair ID
    phi_coefficient = phi_coefficient,
    pair_nesting_pattern = pair_nesting_pattern, # Add nest attendance pattern
    male_nesting_count = male_nesting_count,  # Male's nest attendance count
    female_nesting_count = female_nesting_count,   # Female's nest attendance count
    iteration = iteration  ## Add iteration number
  ))
}


# Run simulation for all pairs ####

# Set number of simulation iterations
num_simulations <- 10000

all_simulation_results <- input_data %>%
  pmap(function(pair_id, male_trip_lengths, female_trip_lengths, observation_period) {
    future_map(1:num_simulations, ~ run_simulation(pair_id, male_trip_lengths, female_trip_lengths,
observation_period, .x),
               .options = furrr_options(seed = TRUE))
  })
```
