## Supplemental material S2 for "Coordinated Discordance: Strategic Nest Attendance for Chick Rearing in Monogamous Seabird"

Distribution of φ coefficients calculated from nest attendance patterns generated by resampling simulations (Supplemental material S1) based on observed trip duration distributions for each pair, and their correspondence with observed φ coefficients. The φ value shown at the top of each graph represents the φ coefficient obtained from the observed attendance pattern, while the percentage in parentheses indicates the percentile of the observed value from the lower bound of the distribution.

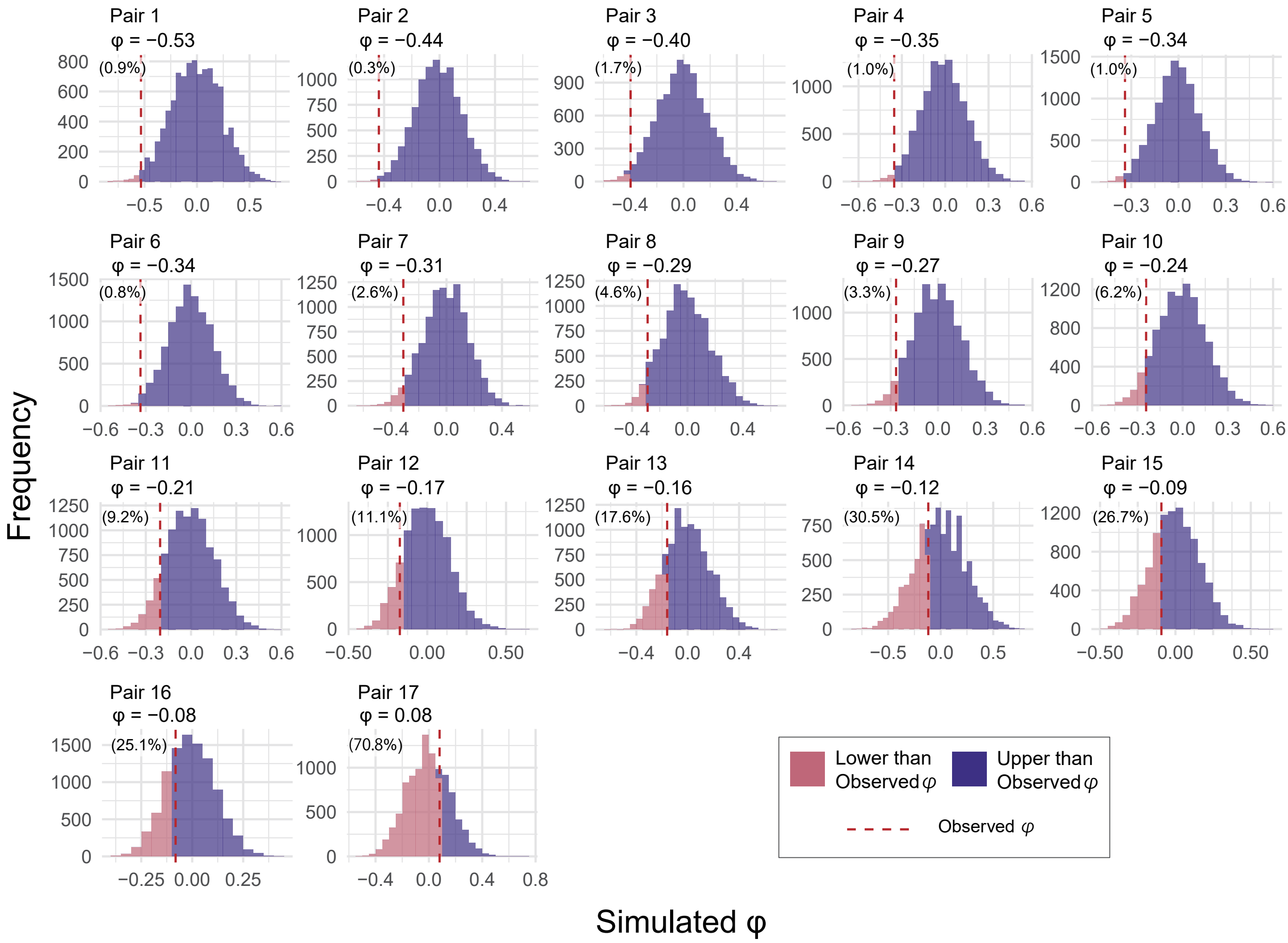
